# Echo-locate: Cerebellar activity predicts vocalization in fruit-eating bats

**DOI:** 10.1101/2024.06.11.598413

**Authors:** Shivani Hariharan, Eugenia González Palomares, Julio C. Hechavarria

## Abstract

Echolocating bats exhibit remarkable auditory behaviors, enabled by adaptations within and outside their auditory system. Yet, research in echolocating bats has focused mostly on brain areas that belong to the classic ascending auditory pathway. This study provides direct evidence linking the cerebellum, an evolutionarily ancient and non-classic auditory structure, to vocalization and hearing. We report that in the fruit-eating bat *Carollia perspicillata*, external sounds can evoke cerebellar responses with latencies below 20 ms. Such fast responses are indicative of early inputs to the bat cerebellum. In vocalizing bats, distinct spike train patterns allow the prediction with over 85% accuracy of the sound they are about to produce, or have just produced, i.e., communication calls or echolocation pulses. Taken together, our findings provide evidence of specializations for vocalization and hearing in the cerebellum of an auditory specialist.

**Teaser:** The cerebellum of fruit-eating bats responds to sounds and predicts future and past vocalizations

## Introduction

Vocal communication is an essential element of mammalian behaviour ^1–4^. It is crucial for social dynamics^5^, and survival^6^. The ability to vocalize is essential for effective communication. A range of animals exhibit this characteristic along with humans, including certain bats^7^, birds^8^, cetaceans^9^, elephants^10^, and pinniped species ^11,12^

While the cortical and subcortical regions involved in hearing^13–16^ and vocalization^17–19^ have been extensively studied, the contribution of evolutionary ancient areas, such as the cerebellum, to vocal behaviour remains an unexplored frontier. In the cerebellum, responses to pure tones or clicks have been recorded from the vermis and hemispheres of passively listening cats ^20,21^, monkeys ^22^ and insect-eating bats^23^. Cerebellar auditory neurons respond to external sounds with fast latencies (typically below 20 ms)^23–25^ and they are thought to receive inputs from classic auditory areas including the dorsal cochlear nucleus at the very gate of the auditory system, the inferior colliculus and auditory cortex^26–29^. Though it is generally accepted that the cerebellum responds to sound, the role of this ancient motor-coordination center in vocalization remains elusive.

In this article, we search for neural correlates of vocalization in the cerebellum of the fruit eating bat, *Carollia perspicillata*. Fruit-eating bats, known for their rich and diverse vocalizations, serve as an excellent model for investigating vocal behaviours in mammals^30–33^. Since the cerebellum of fruit-eating echolocating bats had not been studied before, we first searched for cerebellar neurons responsive to sound, similar to those reported in other mammals. We found clear spiking and local field potential (LFP) responses to sound in the cerebellum of fruit-eating bats, with average latencies ∼ 15 ms in awake and anesthetized preparations. After establishing fruit-eating bats as a good model to study cerebellar auditory responses, we searched for a neural correlate of vocal production within this brain region. We investigated spike trains and LFPs occurring before and after vocalization and found that the type of sound produced (echolocation or communication) can be decoded from pre-vocal and post-vocal neural signals, with prediction accuracies that reach above 85%. The latter provides a direct correlate of vocalization in an ancient motor-coordination structure that lies outside of the classic ascending auditory pathway.

## Results

In this study, we recorded spiking activity and LFPs in the cerebellum of seven adult short-tailed fruit bats (*Carollia perspicillata*), comprising three males and four females, using linear electrode arrays. In the first step, neural data were obtained in anaesthetized and awake bats subjected to passive auditory stimulation, focusing on pure tone stimuli to construct frequency/level receptive fields. We investigated sound-related potentials and spike-sorted single unit responses to acoustic stimuli in various regions in the cerebellum, including the vermis (VIP, VIIa, VIIp, and VIII) ^23,34–36^ and hemispheres (crus I and II) ^20,37,38^, under both anaesthetized and awake conditions. All recordings were conducted on the left-brain side.

Across all multichannel electrode penetrations, sound-evoked responses within the cerebellum were consistently observed. Neural activity was broadly evoked by most frequencies tested and strongest at sound pressure levels above 60 dB SPL (**Fig. 1A and D**, see also individual example of receptive fields in **S1**) Broad, frequency unselective responses were observed whether receptive fields were computed from spike counts evoked by the different combinations of frequency and levels tested (**Fig. 1A**), or by measuring the peak-to peak amplitude of event-related potentials (**Fig. 1D**).

**Fig. 1.**
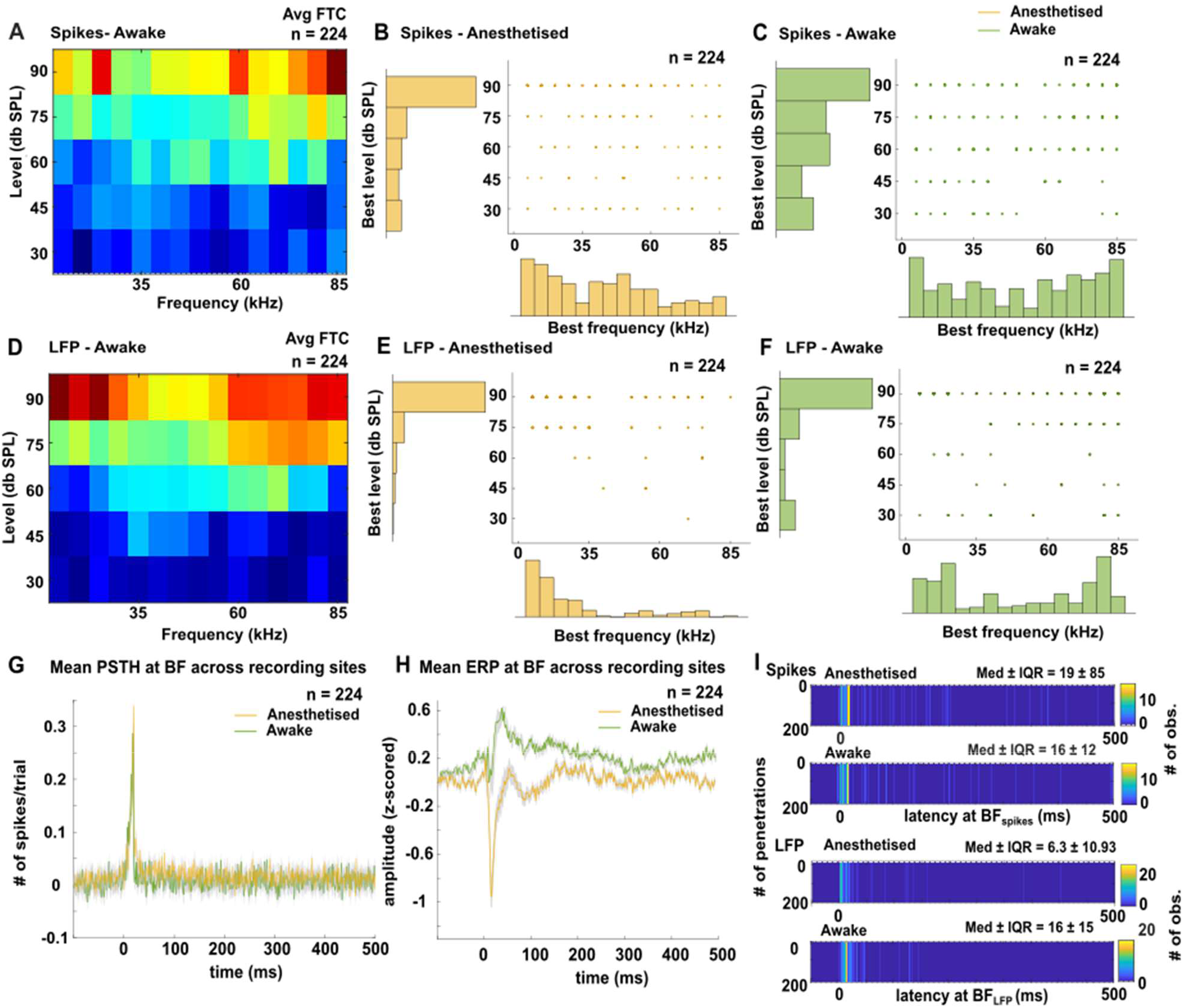
Response to Sound in Anaesthetized and Awake Bats. **A, D**: Average frequency tuning curve (FTC) of 224 recording sites obtained with spike counts (**A**) and the peak-to-peak amplitude of evoked LFP signals (**D**). **B, F**: Distributions of Best Frequencies and Best Levels for the spiking (**B, C**) and LFP signals (**E, F**). Note that in the receptive field, the best frequency/best level combination denotes the point of strongest response. **G, F**: Mean peri-stimulus time histogram and event-related potentials, respectively, obtained at the best frequency/best level combination in awake and anesthetized bats (mean ± SD). **I:** Distributions of latencies measured from spiking and LFP responses.

We computed the best frequency (BF) and best level (BL) of each receptive field, depicting the sound frequency and level that triggered the strongest responses in each unit. In both awake and anesthetized preparations, there was a clear preference for louder sounds (>60 dB SPL) of either low (< 35 kHz) or high frequencies (>60 kHz). Preference for louder sounds is typical of auditory neurons especially in lower stations of the ascending auditory pathway or neurons that receive projections from those areas ^39,40^. When we analyzed the fine temporal structure of sound evoked responses, we observed that the spiking responses observed at the BF/BL combination were generally short and started only a few milliseconds after sound presentation (**Fig. 1G**). Event-related potential obtained at the BF/BL also appeared to have a short latency (**Fig. 1H**). When latency was quantified at the population level, we observed average latency values of spiking activity ranging from 4 to 20 ms (**Fig. 1I**). Most observed neurons displayed a latency of <20 ms, with median values of 19 ms in anaesthetized bats and 16 ms in awake bats. Notably, this latency response in the cerebellum is as fast as that found in insect-eating bats^23^, highlighting the remarkable speed of cerebellar processing in this animal group regardless of feeding behavior. The cerebellar LFPs also demonstrated exceptionally short latencies of <20 ms, with a median latency interval of 6.31 ms ± 10.93 ms in anaesthetized bats and 16 ± 15 ms in awake bats (median± interquartile range, IQR). Similar median short latencies below 20 ms were observed if other parameters of the frequency receptive fields were considered (see supplementary **Fig. S2** for data collected at the characteristic frequency and minimum threshold). Taken together the data collected in passively listening bats both in awake and anesthetized states indicates prominent and fast responses to sound in the bat cerebellum. The latter indicates that the cerebellum plays a role in processing externally produced sounds. But is the bat cerebellum also involved in processing and coordination of self-produced sounds?

### A neural correlate of self-produced vocalizations in the bat cerebellum

To study cerebellar activity during vocalization we placed individual bats with implanted, head-fixed electrodes in an acoustically and electrically isolated chamber and allowed them to vocalize spontaneously while we measured neural activity in the cerebellum. We considered a total of 226 spontaneously emitted vocalizations for investigation after discarding all “unclean calls”, i.e. vocalizations that were not both preceded and followed by at least 500 ms of acoustic silence. Note that most vocalizations produced by bats occur in bursts. In this dataset, clean vocalizations represented 226/1836 of the recorded calls (12.34%). While producing call sequences is the natural behavior for bats, “unclean” calls do not allow to assess uncontaminated pre-vocal and post-vocal activity needed for searching for neural correlates of vocalization. Similar analysis has been used to study vocalization-related activity in the frontal and auditory cortices of this bat species^18,41^

Bat vocalizations fall into two distinct types^41,42^: echolocation and communication (**Fig. 2A**). Echolocation serves primarily for navigation purposes ^43,44^, while communication calls facilitate interaction with other bats^45–48^. We classified spontaneously emitted sounds into echolocation and communication based on their spectro-temporal structure. Echolocation pulses of *C. perspicillata* are short (<2 ms), downward frequency modulated, and peak at high frequencies (>50 kHz)^49^, while communication calls cover a wider range of sound durations and contain most energy at lower frequencies, generally below 50 kHz ^50,51^.

**Fig. 2.**
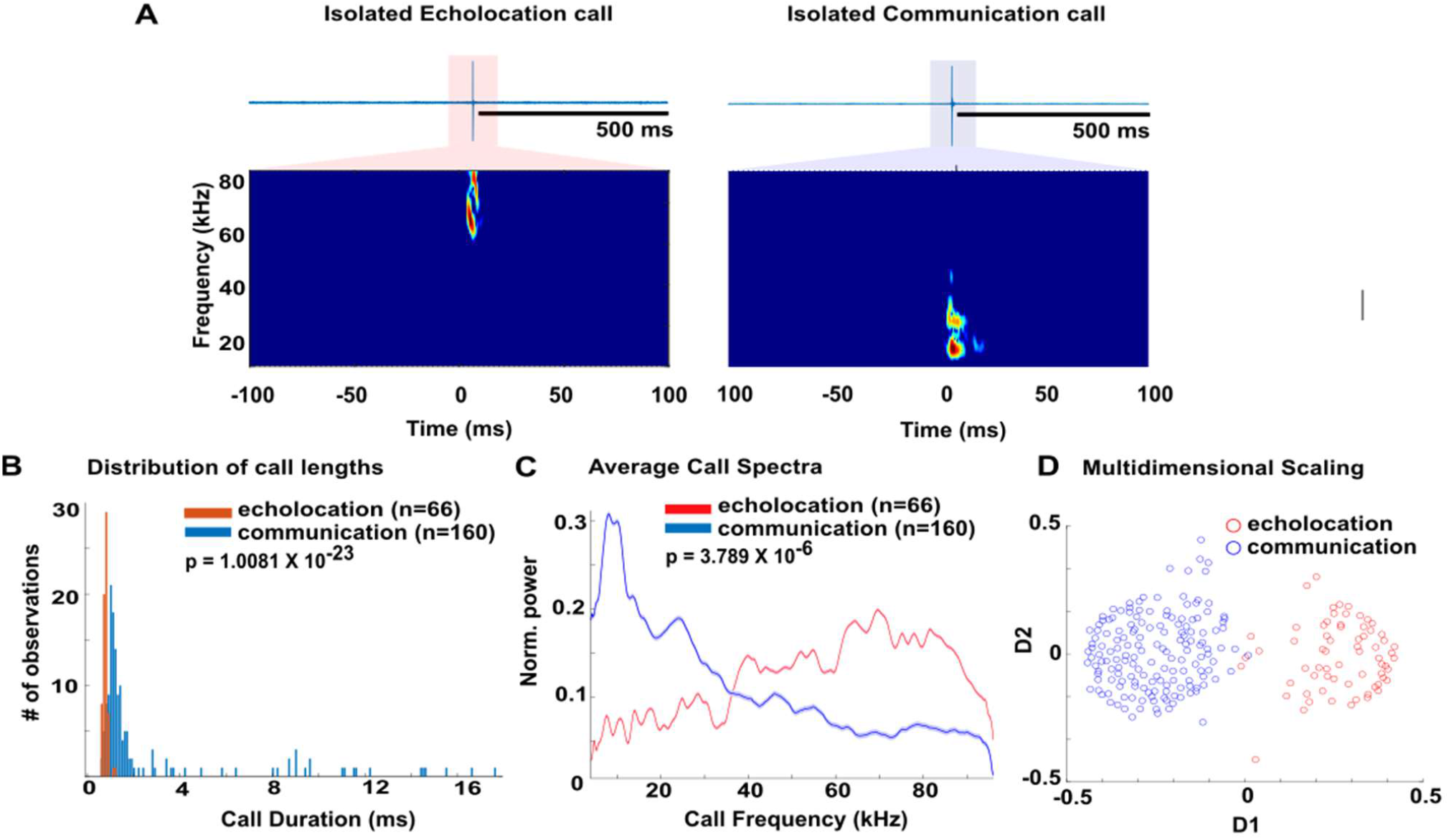
Distinct types of short-tailed fruit bat (Carollia perspicillata) vocalizations. (**A**) Examples of echolocation and communication calls and their respective spectrograms. (**B**) Distribution of call durations. (**C**) Average spectra of echolocation and communication calls (mean ± SD). (**D**) Multidimensionally Scaling of echolocation and communication calls (see Methods for more details).

In our dataset, at the population level, the call duration (**Fig. 2B**) and spectral composition (**Fig. 2C**) of both types of isolated vocalizations differed considerably. Differences between echolocation and communication calls reported here were also evident in multidimensionally scaled data calculated considering the spectral cross-correlation of all calls from each call type (**Fig. 2D**).

After identifying clean vocalizations of each type, echolocation and communication, we explored whether pre-vocal spiking activity in the cerebellum could accurately predict the subsequent vocalization type. Spiking activity occurring 500 ms before and after vocalization onset was analyzed. The average spiking response obtained in the cerebellum for each type of call is shown (**Fig. 3A**). In total, we recorded activity in 1052 and 2535 instances of echolocation and communication production, respectively. These numbers stem from the number of calls produced by the bats multiplied by the number of neurons detected in the multichannel probe (usually 16). Close to the onset of echolocation pulses and communication calls, evident deflections in the spiking signals were observed, potentially indicating evoked responses to the self-produced sounds. Based on the spike shape we could determine that most cells recorded were to Purkinje cells^52–54^ with narrow spikes (217/226, 96.01%, **Fig. 3B**).

**Fig. 3.**
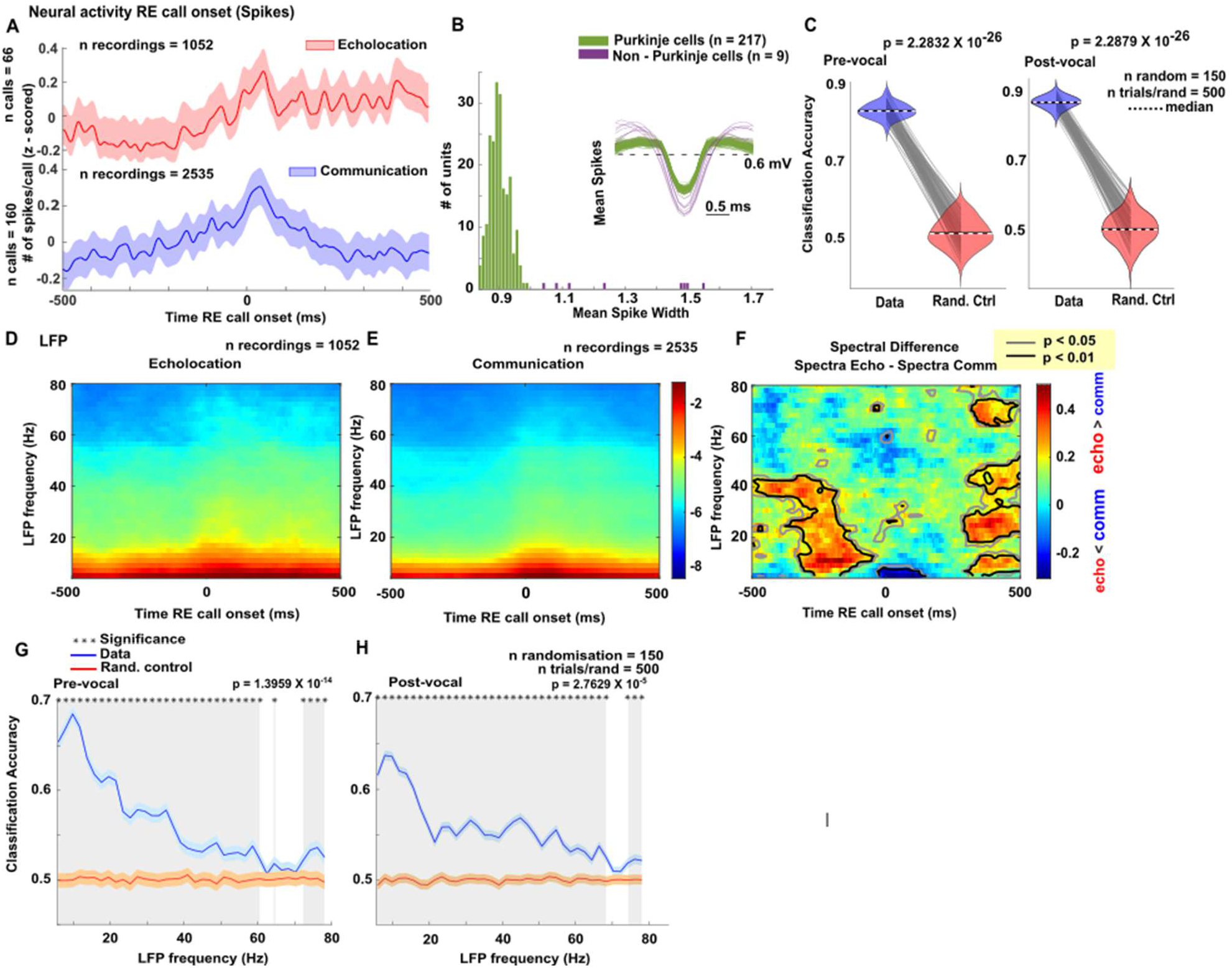
Cerebellar response to vocalizations. (**A**) Spiking neural response during vocalizations relative (RE) to call onset during echolocation and communication (mean ± SD). (**B**) Spike waveforms of non-Purkinje cell type (purple) and Purkinje cells (green). Bottom, Distributions of the mean spike width of corresponding neurons. (**C**) Prediction accuracy calculated using a binary SVM classifier trained with spiking data during vocalizations. Power spectrogram in the cerebellum during (**D**) echolocation and (**E**) communication. (**F**) Spectral difference between echolocation versus communication showing comparisons at each time point and frequency. SVM classifier trained with LFP data occurring (**G**) pre-vocalizations (**H**) post vocalizations onsets (500 ms), (mean ± SD).

To assess whether spiking activity differed in the echolocation and communication conditions, we trained Support Vector Machine (SVM) binary classification models that used the vocalization type (echolocation or communication) as class labels, and the time-binned (bin size = 5 ms) vocalization-related spike trains as predictors. In total, we generated 150 SVM models each taking into consideration the neural signals in 500 randomly chosen vocalization trials (250 echolocation and 250 communication). In each SVM model, half of the trials were used as training set and the other half for blind classification. For each model, we also generated a control condition in which a second model was created in parallel by randomizing the class labels used for training. This allowed direct paired comparison of the models trained with real and randomized data. Our findings demonstrate high vocalization prediction accuracy, achieving 85% and 88% correct blind classification when using pre- and post-vocalization spiking signals, respectively **(Fig. 3C)**. These results highlight the vocalization-specific nature of cerebellar spiking in highly vocal animals, such as bats.

We also performed spectral analysis of the LFP signals related to vocalization. The cerebellar spectrograms of each vocalization type followed the typical power rule by which high power occurred in the low LFP frequencies and power decreased as LFP frequency increased (**Fig. 3D, E**). We then quantified whether the neural spectrograms of LFPs associated with echolocation and communication calls were different (**Fig. 3F)**. For this comparison, we performed FDR^55,56^ corrected Wilcoxon ranksum tests comparing the energy values observed at each time and frequency point in the echolocation and communication-related neural spectrograms (black and grey lines in **Fig. 3F**). We observed several clear spectro-temporal “islands” in which the LFPs associated to echolocation had more spectral power than those linked to communication production (red areas in the spectral difference, blue areas would indicate the opposite trend, i.e. communication > than echolocation). The p-values obtained indicated strong to moderate differences in the neural spectrograms associated to echolocation and communication (p-value<0.05= moderate differences, p-value<0.01= strong differences). Differences in the neural spectrograms generally occurred at ∼ 250 ms before vocalization and in the late periods after vocalization onset (i.e. ∼ 400 ms post-voc) and they were strongest for LFP frequencies below 60Hz in the pre-vocal period but could also reach the high gamma band (i.e., >60 Hz) in the late post-vocal time window. LFP power was typically strongest during echolocation (red areas in **Fig. 3F**), except for the time point of vocalization at which communication-related neural spectrograms has more power in the low LFP band (< 8Hz, dark blue areas in **Fig. 3F**).

In general, these results suggest that, like with spiking, cerebellar field potentials could be used to predict which vocalization the bats are going to emit before they actually do it. This result was corroborated using SVM classifiers trained in the same way as for the spiking data (i.e. 150 SVM models each considering neural signals from 500 randomly chosen vocalization trials). Note that in the case of LFPs, the predictors are the time signals associated to each frequency in the neural spectrogram, and thus we can assess how well individual neural frequencies predict vocal output. The results showed that, indeed, cerebellar field potentials recorded before and after vocalization onset can be used to predict the vocalization type. Predictive power was highest in the low frequencies although it was generally above the baseline randomized control for a broad frequency range. In this sense, it seems that cerebellar LFPs are broader than cortical ones, where vocalizations can be predicted from narrow frequency bands in the LFP^18,41^ Interestingly, predictive power from cerebellar LFPs reached a maximum of 68% which was lower than that obtained from spiking signals (maximum of 88%, see above), suggesting that in the cerebellum, spiking signals are a better read out of vocal behavior when compared to LFPs.

## Discussion

The role of the cerebellum in vocalization remains relatively underexplored compared to the extensively studied cortical and subcortical regions involved in auditory and vocal processing. This study provides evidence of cerebellar responses to auditory stimuli and vocalization in fruit-eating bat, *C. perspicillata*. Our findings reveal that the cerebellum exhibits rapid responses to auditory stimuli and its activity provides a neural correlate of vocal production. Future, ongoing and past vocal behaviors can be predicted from cerebellar spiking and field potentials. Our data indicates that the mammalian cerebellum participates in both vocal coordination and hearing.

The main contributions of the present article can be split into the roles of the cerebellum in 1) hearing and 2) vocal behavior. In terms of hearing, this paper demonstrates that the bat cerebellum responds to external sound stimuli as simple as pure tones. Auditory responses in the bat cerebellum had been characterized before in insect-eating bats that specialize in hunting moving prey using their sonar^23,36^. Fruit eating bats, like the one studied here, do not hunt moving prey but still use their sonar for general orientation^49,57,58^. Our results show that, much like insect-eating bats, the cerebellum of frugivorous bats contains auditory neurons that respond robustly to sound with short latencies typically ranging from 4-20 ms. Auditory functions in the bat cerebellum could have been already present in an ancestor common to both extant insectivorous and frugivorous bats. The function of the cerebellum in complex computations can be tracked down in the evolution to species such as fish which perform electrolocation^59–61^. Bats are in fact not the only mammalian clade in which auditory responses in the cerebellum have been found. This list includes rats^1^, cats^20,21,62^, and monkeys^20–22,63^. Also in humans, the cerebellum seems to play a role in vocalization planning and perception^37,38^. Our data obtained in bats supports this notion. The bat cerebellum with its rapid, phasic neuronal responses to sound could be involved in auditory perception.

A clear finding of our study is the predictive power of the cerebellum in the context of vocal communication. Using pre-vocal spiking measured in the bat cerebellum, we could predict with >80% accuracy the type of calls bats were about to emit. Such high predictive power was only found before in the frontal cortex of the same bat species when using extensively averaged LFPs^41^. In our data, cerebellar LFPs rendered lower predictions than cerebellar spiking. A possible difference between the predictive power of two types of neural signals could be related to the diversity of neurons and physical structure in the cerebellum and how these contribute to summed population signals measured in LFPs. Cerebellar activity could also predict vocalization type in the post-vocal periods, highlighting a role of the cerebellum beyond vocal coordination and in agreement with our results obtained in passively listening animals.

Taken together, the findings depicted in this article demonstrate that the cerebellum of an auditory specialist, a fruit eating bat, responds to external sounds in a fast and robust manner. Cerebellar activity contains a strong correlate of future, ongoing and past vocalizations suggesting a role of this ancient structure in vocalization and hearing. It seems that, though evolutionarily ancient, the cerebellum is still part of the acoustic communication brain network in extant mammals.

## Methods

In this study, 7 adult animals (3 males and 4 females; mean initial weight ± standard deviation (STD): 22.7 ± 3.47g and 19.2 ± 0.15 g respectively) of the bat species *Carollia perspicillata* were used. These bats were sourced from the colony at the Institute for Cell Biology and Neuroscience, Goethe University, Frankfurt am Main, Germany. The animals were housed in a temperature-controlled room maintained at 28°C with a humidity level of approximately 60%. Our experiments adhered to current German laws governing animal experimentation (*Experimental permit No. FU-1126, Regierungspräsidium Darmstadt*).

### Experimental Design

#### Surgical procedure

Before surgery, bats were anaesthetised subcutaneously with a mixture of Ketamine (*10 mg/kg Ketavet, Pfizer, Berlin, Germany*), Xylazine (*38 mg/kg Rompun, Bayer, Leverkusen, Germany*) and Sodium chloride in a ratio of 20:2:18 respectively. Local anaesthesia (*Ropivacaine 1%, AstraZeneca GmbH, Wedel, Germany*) was administered topically. The animals were used for a maximum of 14 days and euthanised with an anaesthetic overdose of pentobarbital (*160 mg/ml Narcoren, Boehringer Ingelheim Vetmedica GmbH, Germany*). The initial dose and subsequent doses administered during the recording period were adjusted for each animal depending on its body weight and resistance to the chemical.

A rostro caudal midline incision was cut, after which muscle and skin tissues were carefully removed to expose the skull. A metal rod (ca. 1 cm length, 0.1 cm diameter) was attached to the bone to guarantee head fixation during electrophysiological recordings. The cerebellum was located through well-described landmarks, including prominent blood vessel patterns^64^. Subsequently, two openings were made into the exposed skull, one for the active derivation of neuronal responses and one to function as a reference for measuring neuronal activity. The experimental animal was allowed resting periods of 3 days after each craniotomy before further experiments were performed. No experiments on a single animal lasted longer than 4 h per day. Water was given to the bats every 1–1.5 h period, and experiments were halted for the day if the animal showed any sign of discomfort.

#### Acoustic Stimulation

During the recording sessions, the neural responses were recorded during two different behavioural contexts: passive listening to auditory stimuli and voluntary vocalizations. In the first context, the bats were presented with pure tones at different frequencies. In the second context, the bats vocalized spontaneously while their vocal output was continuously monitored with a microphone placed at mouth level (10 cm distance). The neuronal activity was recorded simultaneously with the vocal output to study the relationship between neural activity and vocal behaviour.

To assess the frequency tuning of the cerebellum, pure tones (10 ms, 0.5 ms rise/fall time) at a frequency from 15 to 85 kHz in steps of 5 kHz with a combination of levels from 30 to 90 dB SPL in steps of 15 dB SPL were played. Thus, there were a total of 15 pure tone stimuli that were played in a pseudo-random total of 8 times, with a pre-time of 250 ms and a post-time of 50 ms. The sound was generated with a sound card (RME Fireface UC; 16 bit precision, 192 kHz; RME Audio, Haimhausen, Germany) at a sampling rate of 192 kHz, then to an audio amplifier (Rotel power amplifier, model RB-1050) and was played through a speaker (NeoCD 1.0 Ribbon Tweeter; Fountek Electronics, China) placed 30 cm away bilateral to the ears. The speaker was calibrated using a microphone (*MK 301, MG Electronics, USA; gain 10V/Pa*) recorded at 16-bit and 384 kHz of sampling frequency with a microphone amplifier (*Nexus 2690, Brüel & Kjær*). The root mean square (RMS) level of the calls used as stimuli was also calibrated to dB SPL using a 1 kHz pure tone at 94 dB SPL as a reference.

#### Classification of calls

Given the stereotyped spectral properties of *C. perspicillata’s* echolocation calls, a preliminary classification between echolocation and communication utterances was done based on each call’s peak frequency (a peak frequency > 50 kHz suggested an echolocation vocalization, whereas a peak frequency below 50 kHz suggested a communication call) ^50,51^. In addition, vocalizations were labelled as candidates for subsequent analyses if there was a time of silence no shorter than 500 ms before and after call production to ensure no acoustic contamination on the pre-vocal and post-vocal periods that could affect LFP measurements in the cerebellum. Finally, echolocation and communication candidate vocalizations were individually and thoroughly examined via visual inspection to validate their classification (echolocation or communication), the absence of acoustic contamination in the 500 ms prior and 500 ms post vocal onset, and the correctness of their start and end time stamps. According to the above, 66 echolocation and 160 communication calls were then used in further analyses.

### Statistical analysis

#### Analysis of LFP and Spike data

To evaluate the frequency tuning of each unit and electrode channel, the response to pure tones from (15 to 85 kHz at 5 kHz steps) and SPL level (from 30 to 90 dB SPL at 15 dB SPL steps) were analyzed. All recorded spike and LFP data were processed with custom MATLAB (2021b) scripts. In the passively listening animal, to characterize the frequency tuning of recording sites, the best frequency (BF, the stimulus frequency that elicits the largest magnitude response ^65,66^) and best level (BL, the stimulus SPL level that elicits the largest magnitude response^67,68^) for a particular recording site was determined by examining spikes and LFPs evoked by the presented frequency-level stimulus set. The BFBL was determined quantitatively by either identifying the frequency and level with the highest spike count in the spiking activity or with the highest instantaneous energy in the LFP response. Population spiking activity was shown as average PSTH in response to BFBL tones.

Neural signals were filtered in the 300-3000 Hz frequency range using a second-order Butterworth filter to observe the spiking activity. The spike waveforms were sorted using an automatic clustering algorithm, KlustaKwik^69^, which uses results from principal component analysis to create spike clusters ^70^. For each recording, we considered only the spike cluster with the highest number of spikes. From the firing of action potentials, peri stimulus histograms (PSTH, 1 ms bin size) were computed.

For LFP analysis, the electrophysiological signal was down-sampled from 20 kHz to 1 kHz, the line noise was eliminated using the Chronux toolbox's rmlinesmovingwinc function^71^, and the electrophysiological signal was filtered between 1 and 80 Hz (second-order Butterworth filter) to study LFPs during each condition. In addition, signals were normalized, and Z scored individually in each channel of the cerebellum. Neural spectrograms of LFPs corresponding to the communication or echolocation condition were obtained using the mtspecgramc function of Chronux ^71^ (window size of 250 ms, 0.5-ms time step, and a time-bandwidth product of 2 with 2 tapers).

Statistical power was evaluated by characterizing the difference between the spectra of echolocation to the spectra of communication related LFPs using Wilcoxon rank sum tests. The Wilcoxon rank sum test compares two populations when samples are independent by returning the rank sum of the first sample. The pvalues of the data were then corrected using the False Discovery Rate (FDR), (function “mafdr”, MATLAB 2023b) that contains a positive False Discovery Rate (pFDR) for each entry in P-Values^55,56^. The two populations compared in the Wilcoxon ranksum tests refer to spectral power values obtained at specific time/frequency points relative to vocalization onsets in the echolocation and communication conditions.

For prediction of vocal output using the average spectral signal in each field potential band and spiking activity a binary SVM classifier either before or after vocalization. Support Vector Machine (SVM) is a discriminative model that separates data into classes by finding the optimal hyperplane that maximally separates the two classes ^72,73^. As a statistical model, SVM can be described in terms of its assumptions, parameters, and performance measures. We used 150 SVM models each considering neural signals from 500 randomly chosen vocalization trials (*fitcsvm function, linear kernel, MATLAB 2021b, single training, standardized*).

## Acknowledgements

We thank Gisa Prange and the Animal caretakers at the Institute for Cell Biology and Neuroscience in Frankfurt for their support. This study was funded by the German Research Council, Grant number 525183217, Grant to JH.

