## Supplementary Figure for "Echo-locate: Cerebellar activity predicts vocalization in fruit-eating bats"

### Supplementary Figures:

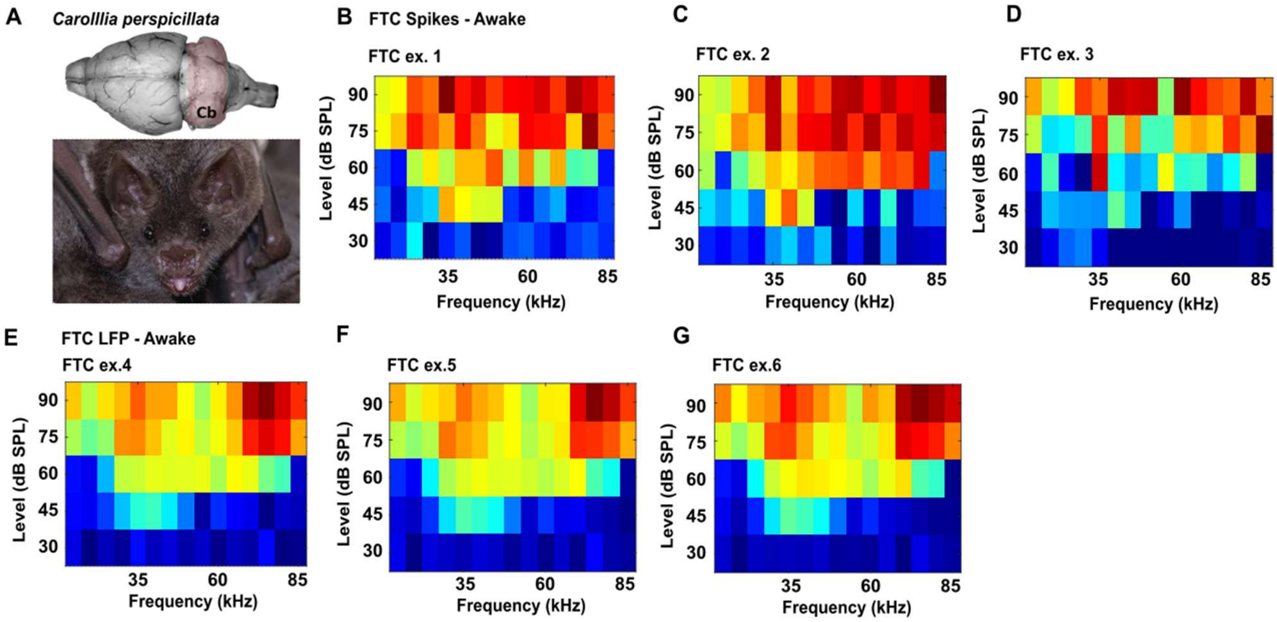


***S1 Examples of FT curve in Anaesthetized and Awake Bats***

(**A**) Picture of bat brain with the area of recording in the cerebellum (top) and a vocalizing bat (bottom). Individual examples of receptive fields obtained with spike counts (**B-D**) and the peak-to-peak amplitude of evoked LFP signals (**E-G**).

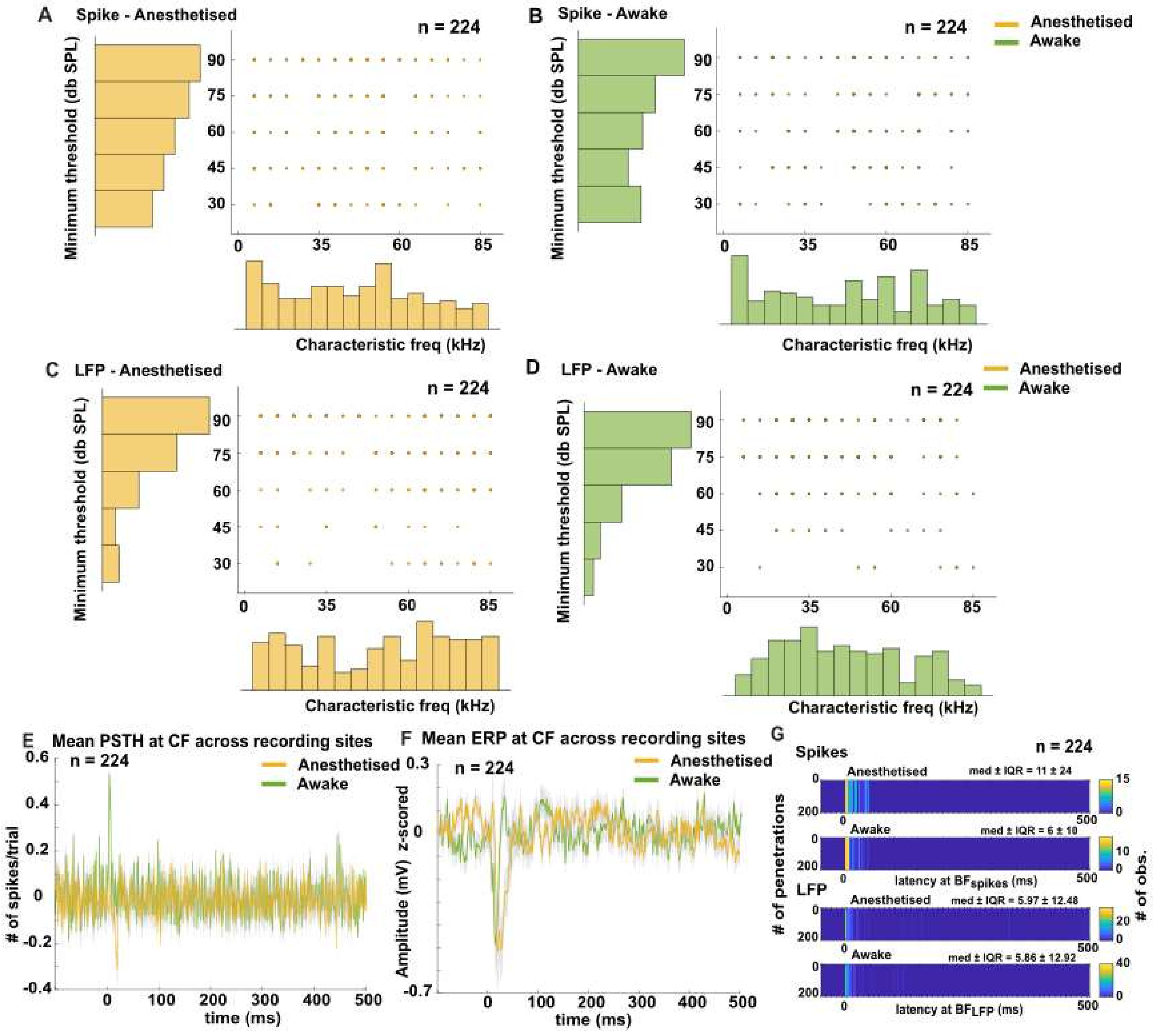


***S2 Characteristic frequency and minimum threshold response in Anaesthetized and Awake Bats***

**A-D**: Distributions of Characteristic Frequencies and Minimum Thresholds for spiking (**A, B**) and LFP signals (**C, D**). Thresholds were determined as the nearest intensity to 65% of the maximum activity observed in each receptive field. Note that the characteristic frequency and minimum threshold combination indicate the point of maximum sensitivity in the frequency/level receptive field. **E, F**: Mean post-stimulus time histogram and event-related potentials, respectively, obtained at the characteristic frequency/minimum threshold combination in awake and anesthetized bats (mean ± SD). **G:** Distributions of latencies measured from spiking and LFP responses.
